# Second-generation method for analysis of chromatin binding using formaldehyde crosslinking kinetics

**DOI:** 10.1101/153353

**Authors:** Hussain Zaidi, Elizabeth A. Hoffman, Savera J. Shetty, Stefan Bekiranov, David T. Auble

## Abstract

Formaldehyde crosslinking underpins many of the most commonly used experimental approaches in the chromatin field, especially in capturing site-specific protein-DNA interactions. Extending such assays to assess the stability and binding kinetics of protein-DNA interactions is more challenging, requiring absolute measurements with a relatively high degree of physical precision. We previously described an experimental framework called CLK, which uses time-dependent formaldehyde crosslinking data to extract chromatin binding kinetic parameters. Many aspects of formaldehyde behavior in cells are unknown or undocumented, however, and could potentially impact analyses of CLK data. Here we report biochemical results that better define the properties of formaldehyde crosslinking in budding yeast cells. These results have the potential to inform interpretations of ‘standard’ chromatin assays including chromatin immunoprecipitation, and the chemical complexity we uncovered resulted in the development of an improved method for measuring binding kinetics using the CLK approach. Optimum conditions included an increased formaldehyde concentration and more robust glycine quench conditions. Notably, we find that formaldehyde crosslinking rates can vary dramatically for different protein-DNA interactions in vivo. Some interactions were crosslinked much faster than the time scale for macromolecular interaction, making them suitable for kinetic analysis. For other interactions, we find the crosslinking reaction occurred on the same time scale or slower than binding dynamics; for these it was in some cases possible to compute the in vivo equilibrium-binding constant but not on- and off-rates for binding. Selected TATA-binding protein-promoter interactions displayed dynamic behavior on the minute to several minutes time scale.

---

Gene regulation is a complicated and highly regulated process involving the coordinated assembly of dozens of proteins on promoter DNA within the context of chromatin (1–4). In vitro studies have provided a structurally detailed paradigm for how the transcription preinitiation complex (PIC) is assembled and regulated (5–13), but less is known about the dynamic assembly of PICs in vivo or how transcription factors (TFs) contribute kinetically to PIC assembly or to the rate of the initiation of synthesis of individual RNAs. To develop molecular models for how these processes occur in vivo, estimates of on- and off-rates for TF binding to specific loci in vivo are required. In instances in which kinetic measurements cannot be made, biophysically rigorous estimates of site-specific in vivo affinity (as opposed to estimates of relative affinity) and fractional occupancy would be valuable.

Chromatin immunoprecipitation (ChIP) is quite possibly the most widely used assay for characterizing the interactions between TFs and specific sites on chromatin. ChIP typically uses formaldehyde to crosslink TFs to their chromatin sites (14), and while it is an undeniably powerful approach for determining transcription factor binding locations with high precision (3), standard ChIP assays are static measurements that do not provide unambiguous insight into the in vivo kinetics of these dynamic interactions. Several assays have expanded ChIP to attempt to capture these relationships. We previously developed a ChIP-based method, the crosslinking kinetics (CLK) assay, which exploits the time dependence of formaldehyde crosslinking to model chromatin-TF binding dynamics on a broad time scale and at individual loci (15). In this approach, cells are incubated with formaldehyde for various periods of time, unreacted formaldehyde is then quenched, and the extent of DNA site crosslinking of a TF of interest at each time point is quantified by ChIP. The time-dependent increase in ChIP signal results from a combination of time-dependent formaldehyde reactivity and time-dependent binding of free TF molecules to unoccupied DNA sites in the cell population. To distinguish kinetic effects of crosslinking chemistry from kinetic effects of TF binding, measurements are made using congenic cells differing only in the concentration of TF and the data are fit using both sets of data simultaneously (15, 16).

A challenge with the development of locus-specific kinetic assays such as CLK is that aspects of the effects of formaldehyde on cells largely remain a black box (17), and validation of the extracted dynamic parameters is difficult because complementary approaches are still being developed and there are few “gold standard” interactions with convergent kinetic measurements obtained by different approaches. Support for the CLK approach was obtained by measurement of binding dynamics for two TFs with very different dynamic properties that had been assessed by live cell imaging (15, 18, 19). However, live cell imaging has its own technical challenges (20) and in most cases it is not possible to identify particular single copy chromatin sites of interaction by live cell imaging (8, 21, 22). An alternative approach is competition ChIP, an assay that measures the rate of turnover between an endogenous and inducible copy of a TF. Our recent work demonstrates that quantitative estimates of locus-specific binding kinetics can be obtained by modeling competition ChIP data, including the estimation of residence times much shorter than the time for full induction of the competitor TF (23). Importantly, comparison of CLK and competition ChIP data for TATA-binding protein (TBP) to a few specific loci shows that the time scales for chromatin interaction are similar as judged by the two methods, with residence times for promoter binding being in general on the order of several minutes (23).

Nonetheless, locus-specific TF-chromatin dynamics are just beginning to be explored, with only a small number of TFs and chromatin sites for which CLK, competition ChIP and/or live cell imaging kinetic data are available. A key aspect of the CLK assay involves the trapping of bound species using formaldehyde. Here we report biochemical results that better define the chemical behavior of formaldehyde in yeast cells. An increased formaldehyde concentration led to more rapid crosslinking, which improved the time resolution and analytical ability of the assay to extract locus-specific binding kinetic information for some TFs. For other TFs, an increased formaldehyde concentration resulted in their depletion from the soluble pool, and in some cases rapid depletion. These observations emphasize the importance of optimizing the CLK approach for analysis of the dynamic behavior of a particular TF. We report the development of a general and improved CLK method framework with both more rapid crosslinking and more efficient quenching in yeast cells. We also report improved computational methods for data analysis and describe improved approaches for distinguishing contributions of crosslinking rate and binding kinetics to the time-dependent increases in ChIP signal.

## RESULTS

The CLK method relies on time-resolved formaldehyde crosslinking ChIP data to assess the kinetics and thermodynamics of TF-chromatin binding. The original CLK method (15, 16) employed 1% formaldehyde (360 mM) and reactions were quenched with 250 mM glycine (24, 25). Under these conditions, the concentration of glycine is sub-stoichiometric to the formaldehyde concentration as added, but crosslinking was performed by adding formaldehyde to cells in YPD medium, which is made from an amino acid-rich extract of yeast cells and as such, the concentration of unreacted formaldehyde that reaches cells under these conditions is unknown and is most likely well below the initial concentration. Order-of-addition experiments showed that 250 mM glycine could block crosslinking of the Gal4-promoter interaction (15), but we noted in subsequent work that quenching may be variably efficient under these conditions (26). Indeed, we have noticed that for unknown reasons the quench efficiency can be variable from experiment to experiment for certain TFs^1^ (27). To better define time-dependent crosslinking behavior and the impact of different quenching conditions on the resulting ChIP signals, data were obtained using 1% (360 mM) formaldehyde and either 250 mM or 2.93 M glycine using the interaction between yeast TBP-myc and the *URA1* promoter as a model interaction. The high concentration of 2.93 M glycine used in this and subsequent experiments was the maximum achievable based on the solubility of glycine in aqueous solution (~3M) and subsequent dilution resulting from addition of a relatively small volume of concentrated yeast cell culture to the quenching solution (see Experimental procedures). For this reason, we refer to this as the “max glycine” quench condition hereafter. As shown in Fig. 1A, the max glycine quench conditions resulted in lower ChIP signals at each time point compared to 250 mM glycine. These results demonstrate that the concentration of glycine used in the quench can have a significant effect on the magnitude of the ChIP signal, suggesting that more robust quenching of formaldehyde can be achieved with a higher concentration of glycine.

**Figure 1.**
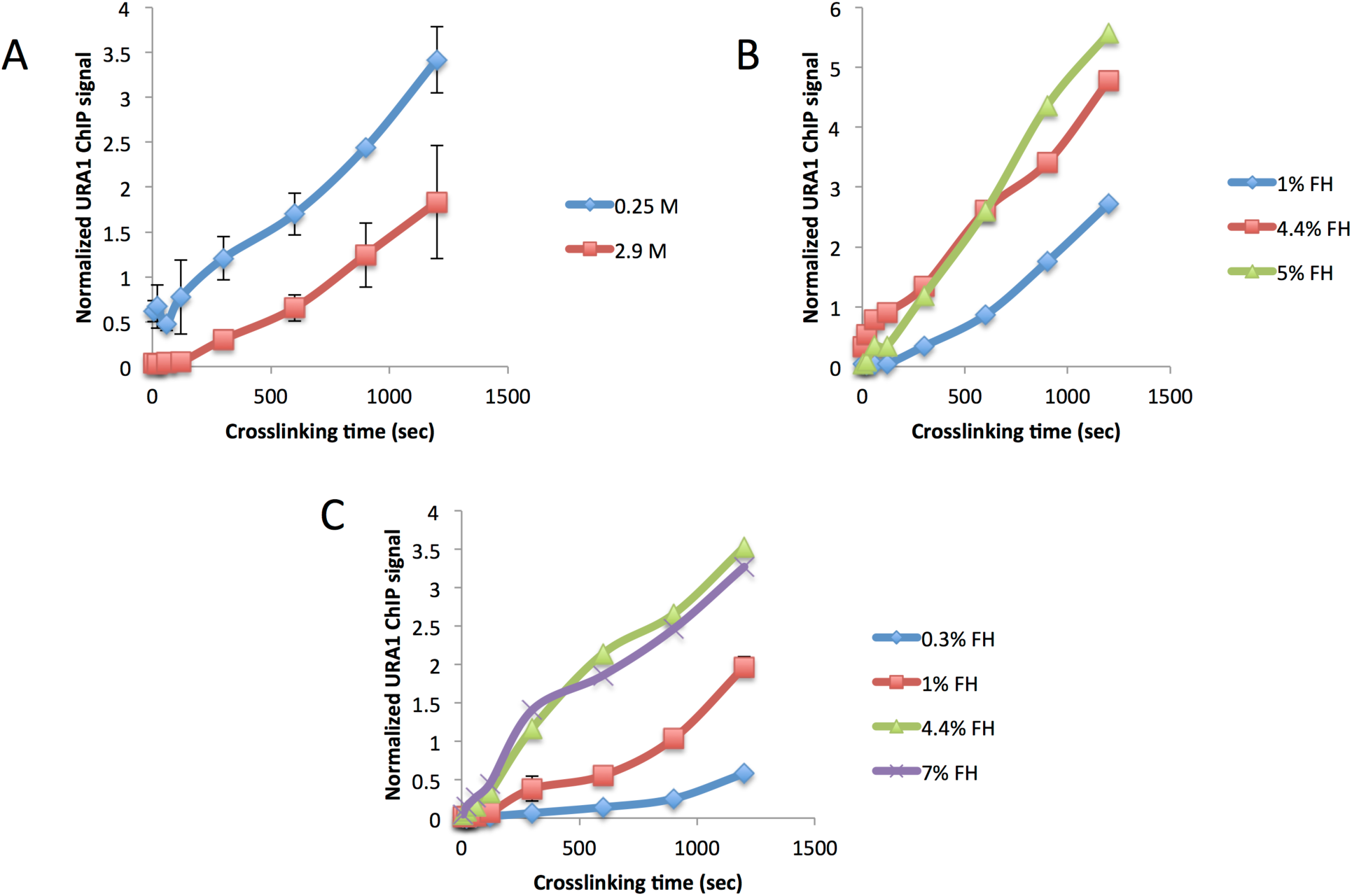
Effect of different formaldehyde and quench conditions on TBP-myc ChIP signal at the *URA1* locus. A) The TBP-myc strain was crosslinked for varying amounts of time with 1% (360mM) formaldehyde followed by quenching with either low (0.25 M, blue line) or high (2.9 M, red line) glycine. Chromatin immunoprecipitation (ChIP) was performed followed by analysis with real time PCR. Normalized ChIP signal is the IP signal minus mock signal divided by an input signal; values were determined from a standard curve. B) TBP-myc cells were crosslinked for varying amounts of time with 1% (blue line), 4.4% (red line), or 5% (blue line) formaldehyde and quenched with high glycine. C) Similar to B, but 0.3% (blue line), 1% (red line), 4% (green line), and 7% (purple line) formaldehyde was used for crosslinking and 600 m Tris pH 8 was used to quench. Error bars represent standard deviation.

In addition to lower signals at each time point obtained using max glycine conditions, some time-dependent datasets showed initial shallow slopes, which continuously increase until the curve reaches apparent linear behavior at longer times (Fig. 1). We refer to this as “positive curvature”. This type of behavior has several possible explanations (discussed below) but none are accounted for in the original CLK model. To better understand how glycine concentration affected the time course of formaldehyde crosslinking, experiments were performed to test both the dependence of the reaction on formaldehyde concentration and how ChIP data were affected using Tris, rather than glycine, to quench the reaction. Tris has been reported to be a robust quencher of formaldehyde reactivity (27). As shown in Fig. 1B, using max glycine quenching conditions, the ChIP signal depended on the formaldehyde concentration, as reaction with 4.4% or 5% formaldehyde increased the ChIP signal at each time point compared to reactions that employed 1% formaldehyde. A dependence on formaldehyde concentration was also seen in reactions using Tris as the quenching agent (Fig. 1C). However, in reactions that were quenched with Tris, the ChIP signals obtained for a given concentration of formaldehyde were reduced compared to the values obtained using glycine, and the resulting reaction progress curves showed positive curvature similar to reactions quenched with max glycine discussed above.

Although Tris is apparently a more efficient quencher than glycine, it also has the potential to reverse crosslinks (28, 29). Crosslink reversal would be problematic for the CLK assay as it could lead to underestimates of ChIP signal, with potentially large percentage-wise effects on the modest levels of crosslinked material obtained after short crosslinking times. To test the potential for reversal with both Tris and glycine, samples were crosslinked, quenched, and incubated at room temperature for different periods of time in the quenching solution. As shown in Fig. 2A, incubation of cells in Tris-containing solution led to a loss of TBP ChIP signal over time. The greatest effect was observed with 750 mM Tris, but the ChIP signal diminished when cells were incubated in 50 mM Tris as well. In contrast, there was no detectable decrease in TBP ChIP signal over time when crosslinked cells were incubated in max glycine solution (Fig. 2B). Thus, although Tris is a robust quenching agent, we ruled out its use in the assay because it decreased the recovery of crosslinked complexes.

**Figure 2.**
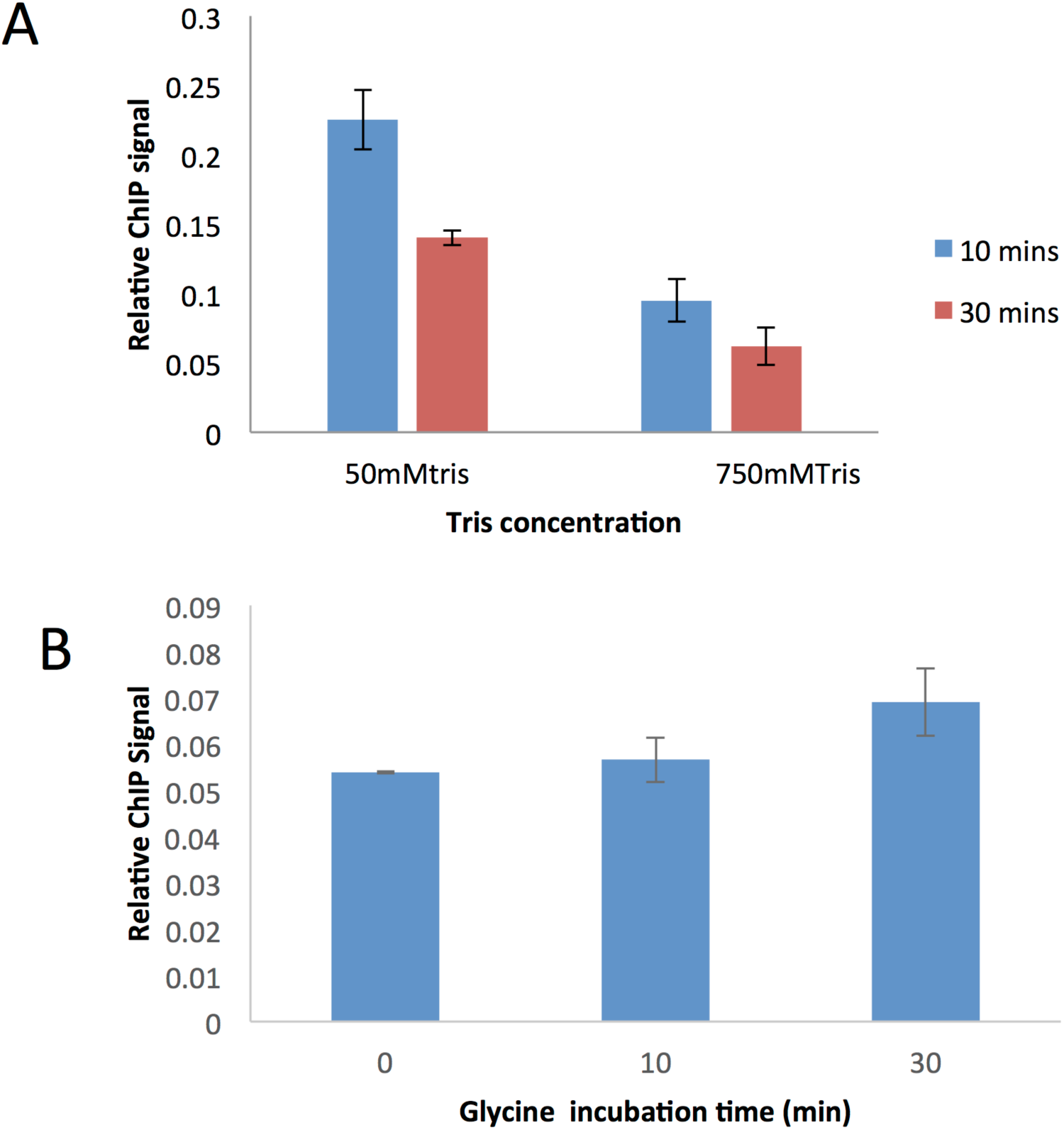
Tris, but not glycine, quenching reverses ChIP signal over time. A) Average ChIP signal of cells crosslinked with 1% formaldehyde, quenched with 250 mM glycine, and resuspended in either 50 mM (blue bars) or 750 mM Tris (red bars), pH 8. Samples were allowed to sit at room temperature for 10 or 30 minutes before processing. B) Average ChIP signal of cells crosslinked with 5% formaldehyde and quenched with 2.93 M glycine pH 5. Samples were left in glycine quench solution for 0, 10 or 30 minutes before processing. All experiments were done in duplicate and error bars are the standard deviation.

The results thus far led to implementation of two significant changes in the CLK methodology. First, to obtain the most accurate time resolved ChIP data, we employed the more robust quenching afforded by max glycine conditions, which lack the negative attributes of Tris as a quencher. Second, as the crosslinking rate is dependent on formaldehyde concentration, we employed 5% formaldehyde rather than 1% as used in previous work (15) (and most ChIP experiments published to date). While 5% formaldehyde optimized the assay for analysis of several interactions in this study, it will be important to determine the optimal formaldehyde concentration for analysis of other types of interactions and in other cell types. We sought the highest feasible formaldehyde concentration for two reasons. First, experimentally, we wanted the ChIP signal to be minimally affected by noise. Second, since the overall crosslinking rate depends on the formaldehyde concentration, faster crosslinking would yield better time resolution between the crosslinking and binding dynamics timescales. To achieve the desired concentrations of reagents in the reactions and to obtain sufficient cellular material for analysis, cell cultures were concentrated by centrifugation, formaldehyde was added to the concentrated cell suspension, and then aliquots of cells were quenched by dilution in a much larger volume of glycine at high concentration. This approach also has the advantage that formaldehyde reactivity is reduced by dilution to 0.1% after glycine addition. Prior work showed that little crosslinking was detectable using 0.1% formaldehyde so dilution alone was expected to have a substantial impact on formaldehyde reactivity (16). In addition, the glycine quenching solution was adjusted to pH 5 which further improves the ability of glycine and formaldehyde to react (27). We refer to the experimental approach employing all of these modifications as CLKv2 (Fig. 3A) to distinguish it from the original CLK method.

**Figure 3.**
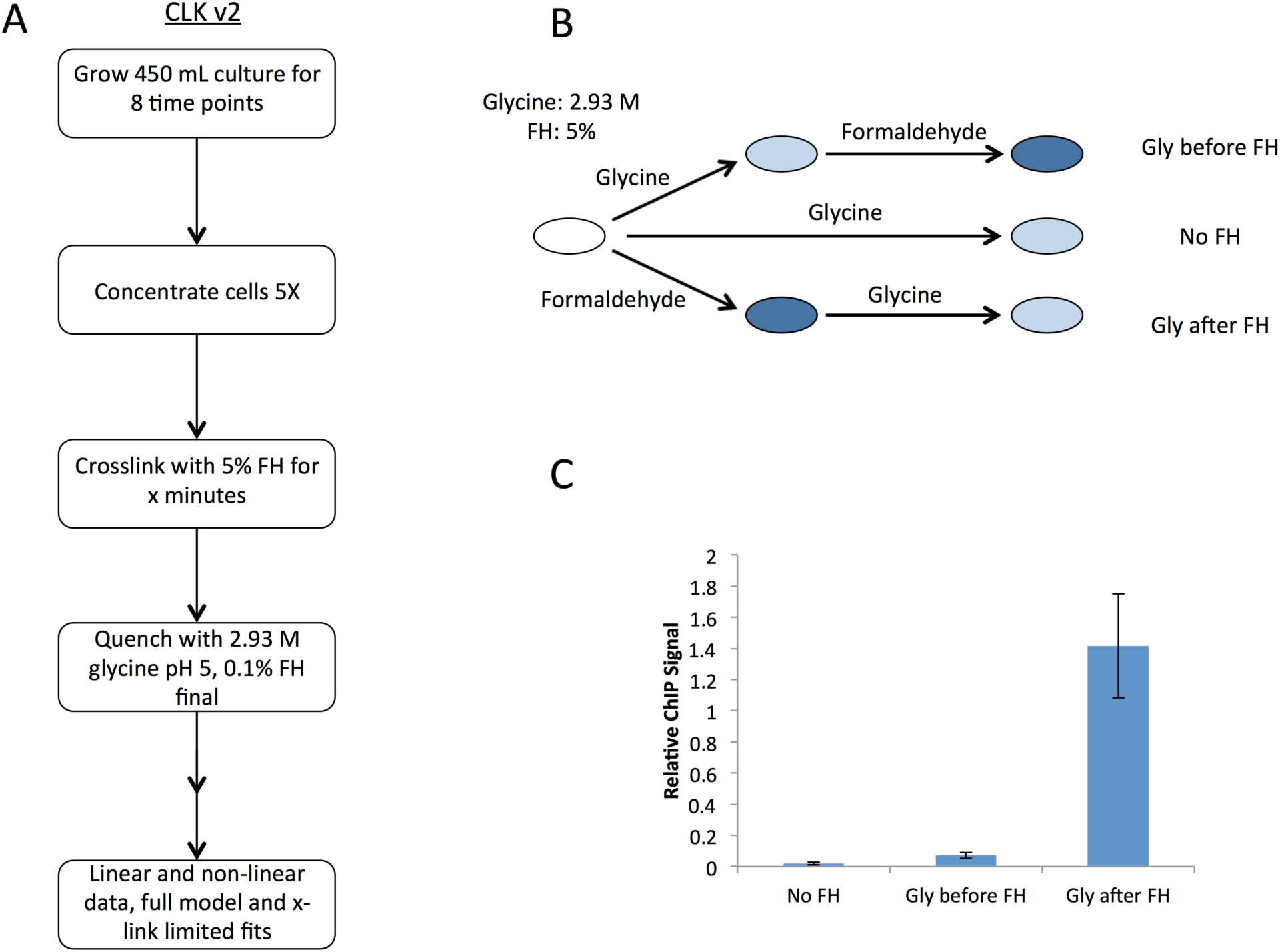
CLK v.2 quenching conditions and overview of the updated method. A) Flowchart of CLK v.2 method focusing on sample collection. B) Order of addition experiments to verify new excess glycine conditions are shown in the schematic. Three experiments were set up: 1) glycine alone added to samples, 2) glycine addition to samples then formaldehyde crosslinking, and 3) formaldehyde crosslinking followed by glycine quenching. For all samples, 5% formaldehyde and 2.93 M glycine pH 5 were used. C) Real time PCR read out from experiments done in B; all conditions were done in duplicate and error is the standard deviation.

As shown in Figs. 3B and C, order-of-addition experiments established that glycine was a very efficient quencher of formaldehyde reactivity when used in this way; the TBP ChIP signal obtained in reactions in which formaldehyde was added first was ~28-fold higher than in reactions with no formaldehyde. In contrast, the ChIP signal obtained when glycine was added before formaldehyde was not statistically different from the background ChIP signal obtained with no formaldehyde at all (p = 0.20). Next, the use of 5% formaldehyde prompted us to evaluate how this higher level of formaldehyde might generally impact cellular constituents. As shown in Fig. 4A, protein yields were reduced in whole cell extracts prepared from cells treated with 5% formaldehyde for increasing periods of time. In contrast, there was no change in the yield of chromatin protein associated with extracts prepared as normally done for ChIP. In addition, there was little change in the pattern of protein bands or their relative intensities over a time course of formaldehyde incubation, indicating that the majority of proteins present in these chromatin extracts were not notably depleted or modified (Fig. 4B). This suggests that the reduced yield of protein in whole cell extracts was due to crosslinked cells being refractory to lysis by rapid agitation with glass beads, whereas soluble protein contents were more efficiently released in the chromatin extract preparation procedure which utilizes a combination of glass bead agitation plus sonication. Protein samples are typically heated to facilitate their denaturation prior to electrophoresis, but formaldehyde crosslinks are also reversible by heat so we analyzed protein extracts on gels with and without heating. There was relatively little difference in overall protein banding pattern when chromatin extract proteins were analyzed following brief heating to facilitate protein denaturation versus unheated samples (Fig. 4B). Heating did reduce an indistinct smear of protein toward the top of the lanes of unheated samples, consistent with heat improving denaturation of the samples. Brief heating had a dramatic effect on the ability to detect TBP in extracts by western blotting (Fig. 4C). The formaldehyde crosslink reversal time is much longer than this brief heating period (30), suggesting that heating in this experiment facilitated disruption of TF-protein complexes and protein unfolding rather than crosslink reversal. In the case of TBP, it is likely that its association with TAFs and potentially other regulatory factors in extracts (31) make detection of monomeric TBP difficult or impossible without heating.

**Figure 4.**
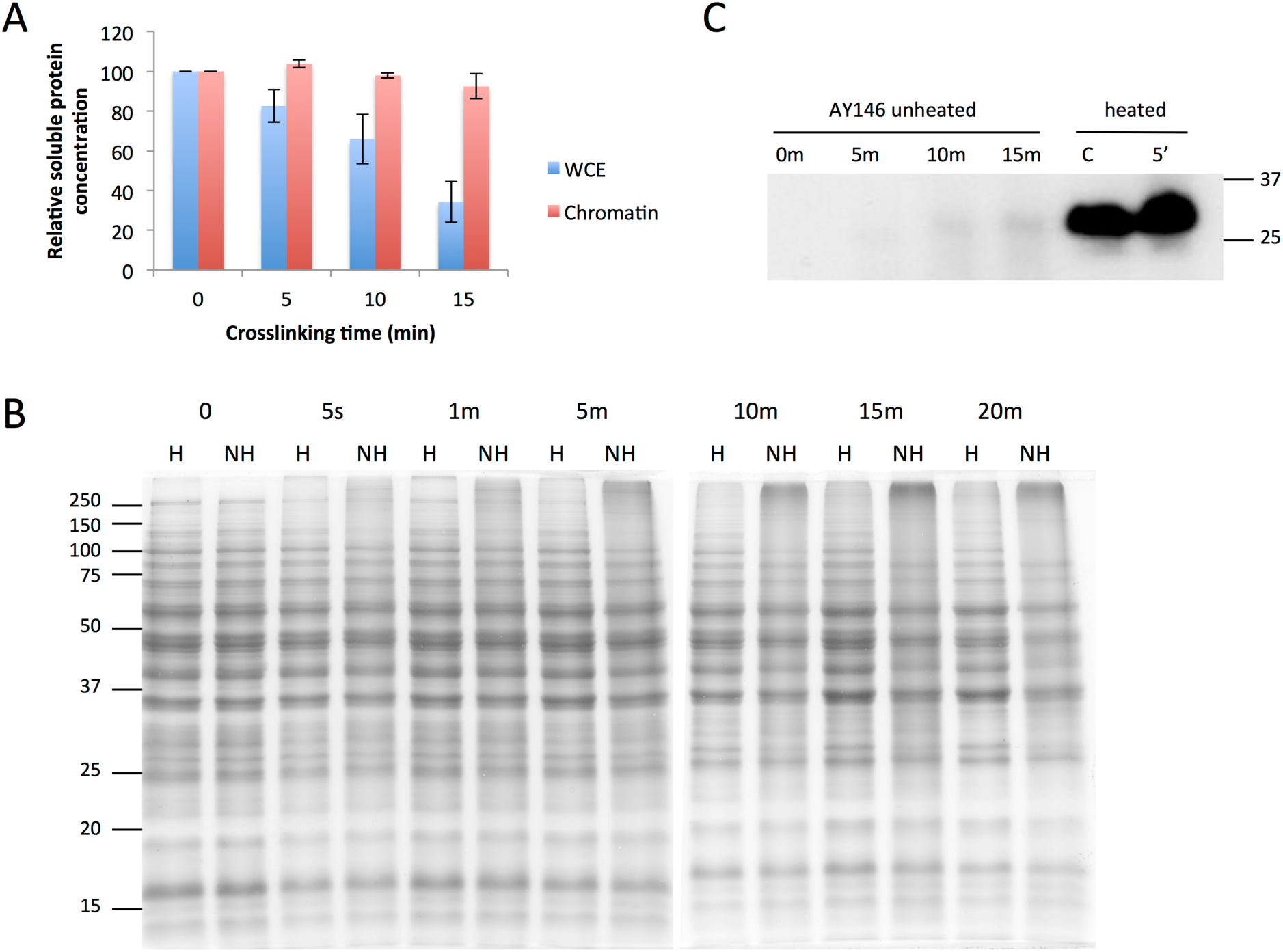
Effect of formaldehyde crosslinking on proteins. A) Relative concentration of protein in either whole cell extract (blue bars) or chromatin (red bars) samples crosslinked with 5% formaldehyde for varying amounts of time. Bradford assays were used to determine the concentration. Duplicates were used for each time point and samples were normalized to their respective zero time point. B) Coomassie stained SDS-PAGE gel of AY146 whole cell extract samples from cells crosslinked for varying amounts of time with 5% formaldehyde. Fifteen microgram samples were heated (H) for 5 minutes at 95°C or not heated (NH) before loading. C) Samples from the AY146 strain were crosslinked for 0, 5, 10, or 15 minutes with 5% formaldehyde and either heated for five minutes at 95°C or unheated before loading into an SDS-PAGE gel. The western was probed with a TBP antibody and visualized with chemiluminescence. C (control) is recombinant TBP protein.

A key requirement for the CLK method is that the unbound pool of the TF being investigated is not depleted significantly by formaldehyde incubation (15). This ensures that there are sufficient molecules available for interaction with unbound DNA sites and that the overall on-rate, which depends on the concentration of the free TF, does not change over the course of the reaction. To determine the effect of 5% formaldehyde on the soluble pools of particular TFs, western blots were performed using extracts obtained from cells treated with formaldehyde for various periods of time. Based on the results in Figs. 4B and C, a brief heating step was used prior to loading samples on the gels in order to accurately estimate the relative amount of soluble TF without reversing any crosslinks that had formed. Western blotting showed that 5% formaldehyde treatment resulted in depletion of some TFs and not others, and the rates of depletion among those that were depleted varied significantly (Fig. 5A-F). TBP, Gal4, and Ace1 were not significantly depleted in these experiments, whereas Reb1, Cat8, Abf1, TFIIB and Tfa1 were stable for ~10 min and then were depleted. In contrast, the largest subunit of RNA polymerase II, Rpb1, and the TFIIF subunit Tfg1 were rapidly depleted. This indicates that some factors such as TBP and Gal4 are readily amenable to analysis by CLKv2. As shown below, others such as Tfa1 can be investigated as long as the crosslinking time course is confined to the period in which the levels of the factor are not depleted. Other factors such as Rpb1 and Tfg1 cannot be investigated at present using these conditions. (However, it should be noted that in principle one could incorporate the TF depletion rate into the dynamic model.)

**Figure 5.**
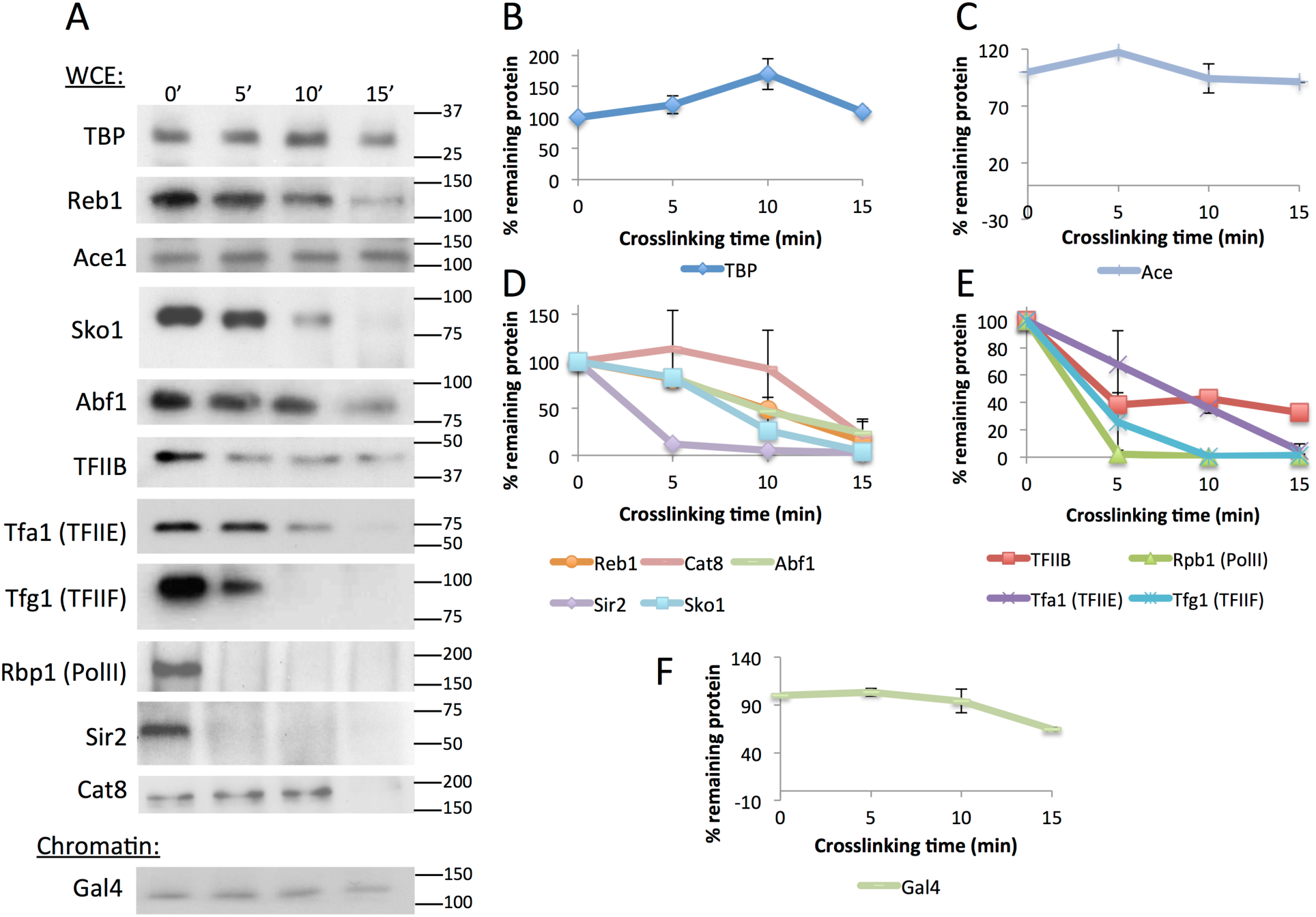
Protein levels in crosslinked whole cell extract or chromatin samples over time. A) Western blots of whole cell extract (WCE) samples for all factors except Gal4, which is a chromatin extract. Antibodies used are listed in Table S4 and molecular weight is denoted to the right in kDa. Samples were crosslinked with 5% formaldehyde for 0-15 minutes and quenched with excess glycine. B) Quantification of WCE western blot bands shown in A for TBP. Each sample was normalized to the 0 time point as a percentage. Two replicates were averaged for the plot and error bars represent standard deviation. C) Same as B, except for Ace1. D) Same as B, except for transcription factors Reb1, Cat8, Abf1, Sir2, and Sko1. E) Same as B, except for preinitiation complex components TFIIB, Tfa1 (TFIIE), Tfg1 (TFIIE), and Rpb1 (RNA polymerase II). Independently performed Western blots using chromatin rather than WCE samples showed the same trends. F) Same as B, except for Gal4 chromatin extract. Gal4 was not abundant enough to be detected in WCEs.

To measure dynamics using the CLKv2 method, ChIP data for an interaction of interest are acquired in two different strains, each of which differ only in the concentration of the TF. One strain (“WT”) expresses the TF of interest at WT levels and the other (“OE” for “over-expression”) typically harbors an additional copy of the TF gene which increases the TF concentration ~2-3- fold on average. The CLK model contains as variables the on-rate for TF-chromatin binding (k_a_), the off rate (k_d_), and the formaldehyde crosslinking rate (k_xl_); the fractional occupancy (θ_b_) and residence time (t_1/2_) are calculated from the variables and are not direct outputs of the fits. The saturation level of the ChIP signal (S_sat_) is an additional parameter obtained from the fits. The concentration of the TF in the nucleus (C_TF_) and the formaldehyde concentration (C_FH_) are experimentally measured quantities used in the fitting calculations. The CLK model makes no assumptions about the relative rates of chromatin binding or crosslinking, and indeed it provides a framework sufficiently flexible to model a wide range of chemical and dynamic behavior (15, 16). Using the CLKv2 conditions, and as discussed in detail below, a wide range of behaviors were observed, including interactions with binding dynamics slower than crosslinking, comparable to crosslinking, or faster than crosslinking. In the binding dynamics-limited scenario (Fig. 6A, D), crosslinking is much faster than the on- and off-rates for chromatin binding. The hallmarks of the binding-dynamics limited behavior (referred to as “TF-limited”) include two exponentials: a very steep exponential rise at short time scales (seconds), often manifesting as a non-zero y-intercept in the WT and OE data with a clear separation in the WT and OE y-intercepts, followed by a slower exponential rise. This clear separation in time scales makes it possible to extract binding dynamics, including the on- and off-rate (15). In contrast, if the rate of crosslinking is slower than the time scale of TF binding dynamics, crosslinking-limited (referred to as “XL-limited”) data show a single exponential rise with a zero y-intercept for the WT and OE data (Fig. 6B). The simulation in Fig. 6B and schematic in Fig. 6E show that for XL-limited interactions, the crosslinking time scale is slower than for the TF-limited case, and under these conditions TF binding and unbinding can occur prior to crosslinking. If the crosslinking rate is so slow (Fig. 6F) that its associated time is longer than the latest crosslinking time (usually 1200 seconds for this study), the ChIP signal rises linearly (or nearly linearly) as shown in Fig. 6C. In the linear version of the XL-limited model the theoretical curve shows a near-zero y-intercept, and no sign of saturation on the experimentally accessible time scale.

**Figure 6.**
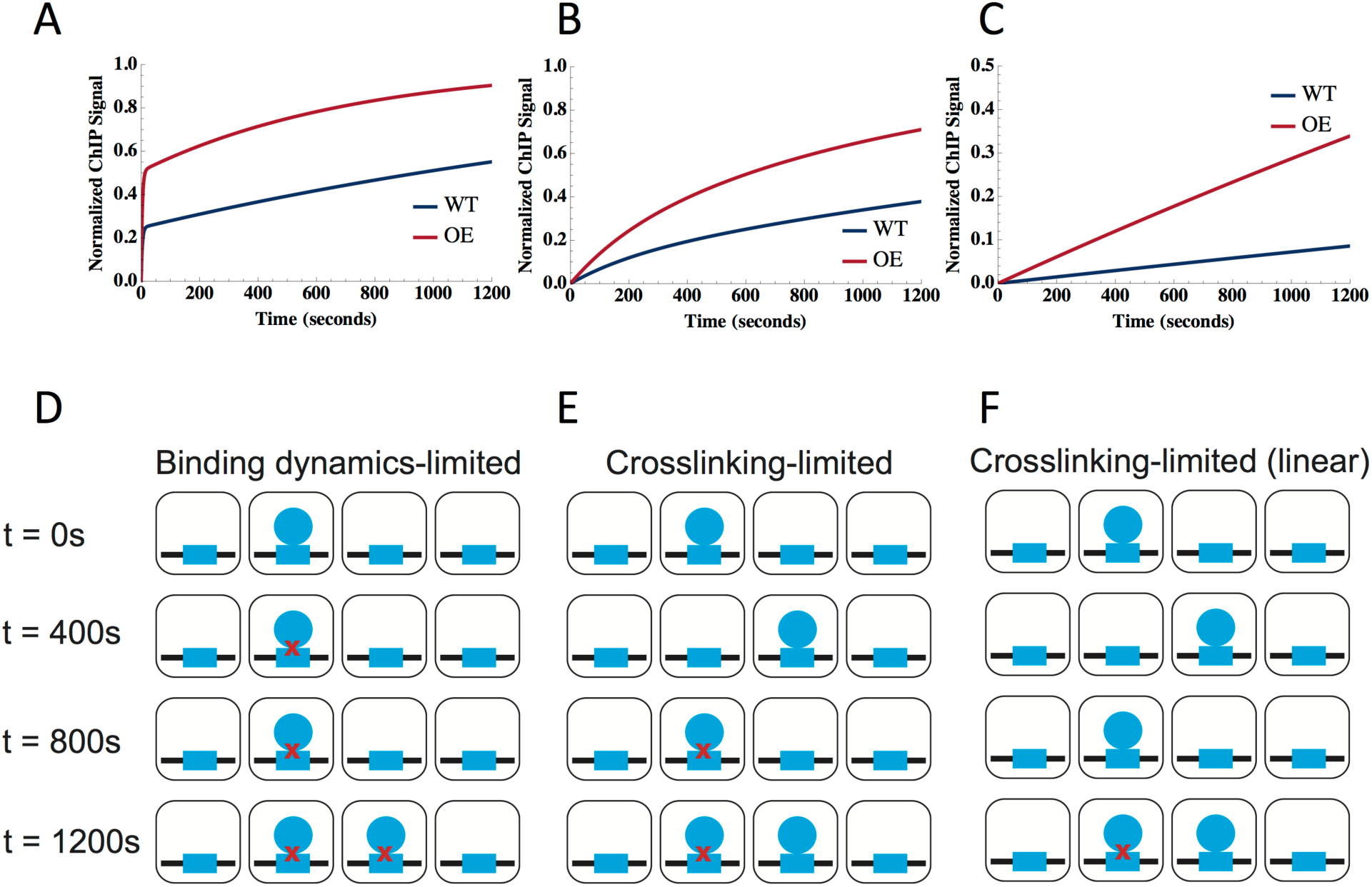
Overview of CLKv2 possible model fits. A-C) Simulations of CLKv2 fits. For each plot, blue represents the wild type strain and red is the overexpression strain. From left to right, the fits correspond to: binding dynamics (TF)-limited (A), crosslinking-limited (B), and linear crosslinking-limited behavior (C). D-F) Schematic of binding dynamics for each of the three CLKv2 cases with formaldehyde crosslinking over time: TF-limited (D), XL-limited (E), and linear XL-limited (F). In each square cell, the TF (blue circles) binds to its binding site (blue rectangles); red x’s represent crosslinking by formaldehyde. Crosslinking time increases as the panels progress from top to bottom.

Once crosslinking time-dependent data have been acquired, determining which scenario describes the data and fitting to the model is described in the flow chart in Fig. 7. The fitting procedures themselves are described in detail in the Methods section. We note that different sets of parameters are gained from each type of fit as shown in the schematic: TF-limited fits yield k_a_, k_d_, k_xl._ and S_sat_ from which the dissociation constant, K_d_, θ_b_, and t_1/2_ can be derived. However, the XL-limited fit only gives K_d_, k_xl_ and S_sat_ from which θ_b_ can be derived and the linear model provides K_d_ and k_xl_*S_sat_ from which θ_b_ can be derived.

**Figure 7.**
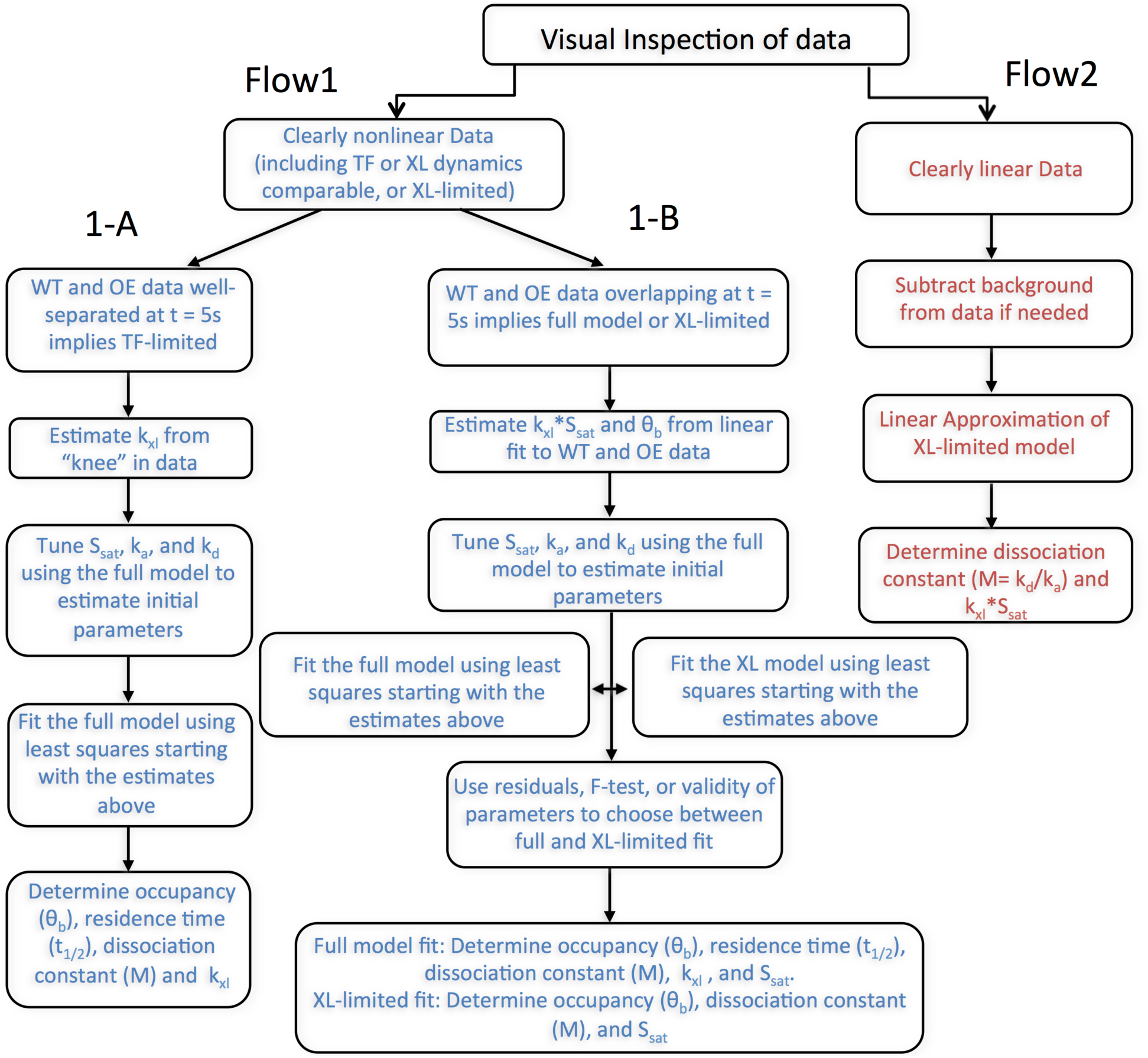
Flow chart for fitting CLKv2 data. After visual inspection of the data, the fitting procedure in either Flow1 arm (blue) or Flow2 arm (red) was followed for a given locus. Fitting procedure for nonlinear data was further broken down into arm 1-A or 1-B. Arm 1-A represents the TF full-model fit, while 1-B could be crosslinking-limited or full model fit. Flow2 is the linear crosslinking-limited fit.

Data were obtained for a number of TF-chromatin interactions using CLKv2. The interactions of TBP with the *LOS1*, *ACT1* and *URA1* promoters are shown in Fig. 8A-C. Applying the flow chart shown in Fig. 7 revealed that these interactions were well described by the TF-limited behavior. At the *URA1* promoter, TBP-myc displayed both a linear ChIP signal with crosslinking time and sensitivity to formaldehyde concentration consistent with XL-limited dynamics, suggesting that although myc-tagged TBP complements growth, the myc tag had a relatively strong effect on crosslinking and possibly TBP binding dynamics. The data describing the interaction between lacI-GFP and an array of lacI sites is shown in Fig. 8D and was also well described by TF-limited behavior. The fractional occupancies of the three TBP loci ranged from 0.04-0.07, while the residence times were about 60-90 seconds (Table 1). Consistent with prior work (15), this indicates that these promoters are unoccupied by TBP most of the time, and that the TBP complexes that do form are not very long-lived. LacI fractional occupancy was lower still, but the complexes formed had half-lives of 1038 seconds. This long lifetime is consistent with both prior CLK and live imaging data (15). TBP binding to *NTS2* (the promoter for Pol I transcription) and Ace1 binding to *CUP1* were both best approximated by the linear model (Fig. 8E, F). The linear behavior of Ace1 CLK data using the CLKv2 conditions is consistent with rapid binding dynamics (15, 32) being faster than the crosslinking rate. The high fractional occupancy of Ace1 at *CUP1* (0.88) is also consistent with prior observations (15, 32). The fractional occupancy of TBP at *NTS2* (0.73) was much higher than TBP occupancies at the other promoters, consistent with the high transcriptional activity of the rDNA in cells in log phase growth in rich medium (33). Table 1 provides all the measured kinetic parameters along with their associated errors. Notably, error analysis derived from multiple fits of simulated data (see Materials and Methods) showed that most parameters were associated with a single well-defined distribution (Figs. S2 and S3)

**Figure 8.**
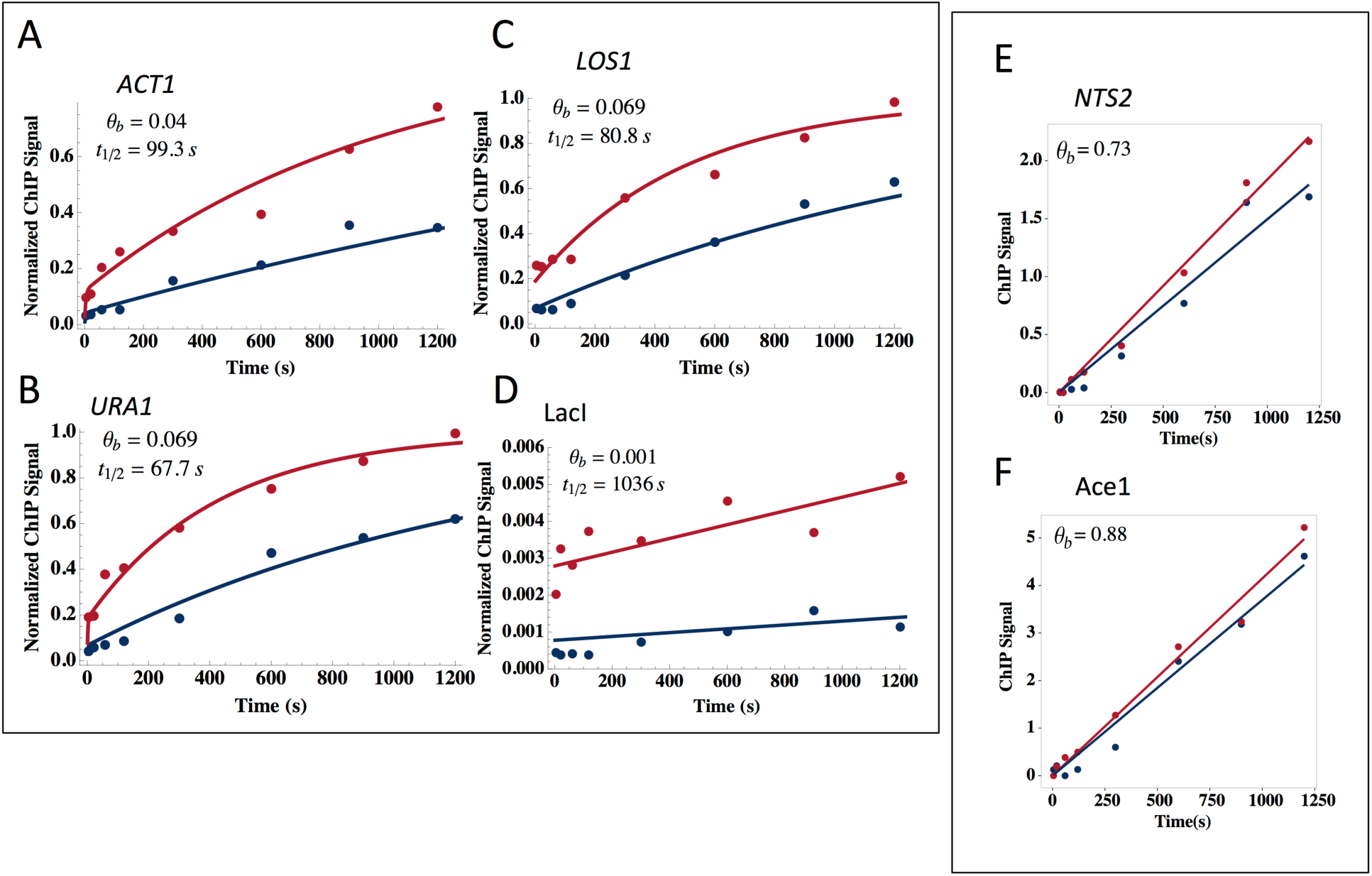
CLKv2 fits of data for TBP, LacI, and Ace1. A-D) All fits shown are TF-limited full model fits. The blue line is the wild type strain, while red has the factor overexpressed; overexpression factors are listed in Table S3. TBP is shown at *ACT1* (A), *URA1* (B), and *LOS1*(C); LacI is shown at a *lac* array (D). Occupancy (0b) and t_1/2_ are denoted on the plots. E-F) Both fits are linear crosslink-limited. TBP is shown at *NTS2* (E) and AceI is shown at *CUP1* (F). Only occupancy (θ_b_) is shown since residence time is not extracted with this fit.

**Table 1.** Measurements for TBP binding dynamics at select promoters.

| TBP-Full model | $k_{xl}$ (1/mol s)<sup>1</sup> | $\tau_{xl}$ (s)<sup>2</sup> | $k_a * C_{TF}$ (1/s)<sup>3</sup> | $k_d$ (1/s)<sup>4</sup> | $S_{sat}$<sup>5</sup> | $K$ (mol)<sup>6</sup> | $t_{1/2}$ (s)<sup>7</sup> | $\theta_b$<sup>8</sup> |
| --- | --- | --- | --- | --- | --- | --- | --- | --- |
| ACT1 | 0.14 (+31.2, -0.51) | 2.76 (+45.05, -0.73) | 3.21 (+1.3, -0.92) E-04 | 6.98 (+3.3, -2.2) E-03 | 1.35 (+0.35, -0.28) | 2.61 (+1.0, -0.74) E-04 | 99.3 (+99.32, -56.51) | 0.044 (+0.44, -0.016) |
| LOS1 | N/A | N/A | 6.31(+1.1, -0.93) E-04 | 8.58 (+2.3, -1.8) E-03 | 0.44 (+0.031, -0.029) | 1.63 (+0.32, -0.27) E-04 | 80.8 (+21.45, -16.94) | 0.069 (+0.012, 0.01) |
| URA1 | 0.30 (+1129.02, -4.74) | 1.29 (+19.22, -0.081) | 7.59 (+1.0, -0.91)E-04 | 1.0 (+0.29, -0.23) E-02 | 0.71 (+0.036, -0.035) | 1.62 (+0.36, -0.29) E-04 | 67.7 (+21.02, -16.2) | 0.069 (+0.015, -0.01) |

| TBP-Linear | $k_{xl} * S_{sat}$ (1/mol s)<sup>1</sup> | $\tau_{xl}$ (s)<sup>2</sup> | $k_a * C_{TF}$ (1/s)<sup>3</sup> | $k_d$ (1/s)<sup>4</sup> | $S_{sat}$<sup>5</sup> | $K$ (mol)<sup>6</sup> | $t_{1/2}$ (s)<sup>7</sup> | $\theta_b$<sup>8</sup> |
| --- | --- | --- | --- | --- | --- | --- | --- | --- |
| NTS2 | 1.1 (+0.084, -0.078) E-03 | N/A | N/A | N/A | N/A | 4.51 (+2.21, -1.44) E-06 | N/A | 0.73 (+0.077, -0.07) |
| SNR6 | 1.7 (+0.12, -0.11) E-03 | N/A | NA | NA | NA | 4.46 (+2.0, -1.37) E-06 | NA | 0.73 (+0.074, -0.067) |
| HSC82 | 3.4 (+0.54, -0.47) E-04 | N/A | NA | NA | NA | 8.96 (+5.73, -3.51) E-06 | NA | 0.57 (+0.13, -0.1) |

| | $k_{xl}$ (1/mol s)<sup>1</sup> | $\tau_{xl}$ (s)<sup>2</sup> | $k_a * C_{TF}$ (1/s)<sup>3</sup> | $k_d$ (1/s)<sup>4</sup> | $S_{sat}$<sup>5</sup> | $K$ (mol)<sup>6</sup> | $t_{1/2}$ (s)<sup>7</sup> | $\theta_b$<sup>8</sup> |
| --- | --- | --- | --- | --- | --- | --- | --- | --- |
| Ace1 (linear) @ CUP1 | 2.34 (+0.1, -0.098) E-03 | N/A | NA | NA | NA | 1.93 (+1.15, -0.69)E-07 | NA | 0.84 (+0.055, -0.052) |
| Lac1 (full model) @ LacO | 1353.9 (+1.21E08, -44332) | 2.84 E-04 (+0.024, -8.68 E-06) | 5.20 (+9.08, -3.12) E-07 | 6.69 (+1.9, -1.5) E-04 | 796.96 (+1421.4, -522.4) | 1.29 (+2.7, -0.9) E-03 | 1037.7 (+328.6, -252.0) | 7.77 (+15, -4.9) E-04 |

<sup>1</sup>Formaldehyde crosslinking rate<sup>2</sup>Crosslinking time<sup>3</sup>On-rate of transcription factor X nuclear concentration of factor<sup>4</sup>Off-rate of transcription factor
N/A: not applicable
<sup>5</sup>ChIP signal at saturation<sup>6</sup>Dissociation constant, $k_d/k_a$ <sup>7</sup>Residence time of TF binding<sup>8</sup>Occupancy

Datasets obtained for TBP binding to the *HSC82* and *SNR6* promoters were not obviously linear or non-linear; these ambiguous cases required a more rigorous selection process for the best fit (see flowchart, Fig. 9A, and Methods section for detailed explanation). These datasets were fit with both the TF-limited and linear models and the sum of squared residuals (SSR) derived from the fits were compared for the appropriate fit (Fig. 9B, C). Both loci had a better fit with the linear model; the SSR for the TF-limited/linear models for *HSC82* and *SNR6* were 0.11/0.042 and 3.35/0.43, respectively. The occupancy of TBP at *HSC82* and *SNR6* was 0.57 and 0.73, respectively.

**Figure 9.**
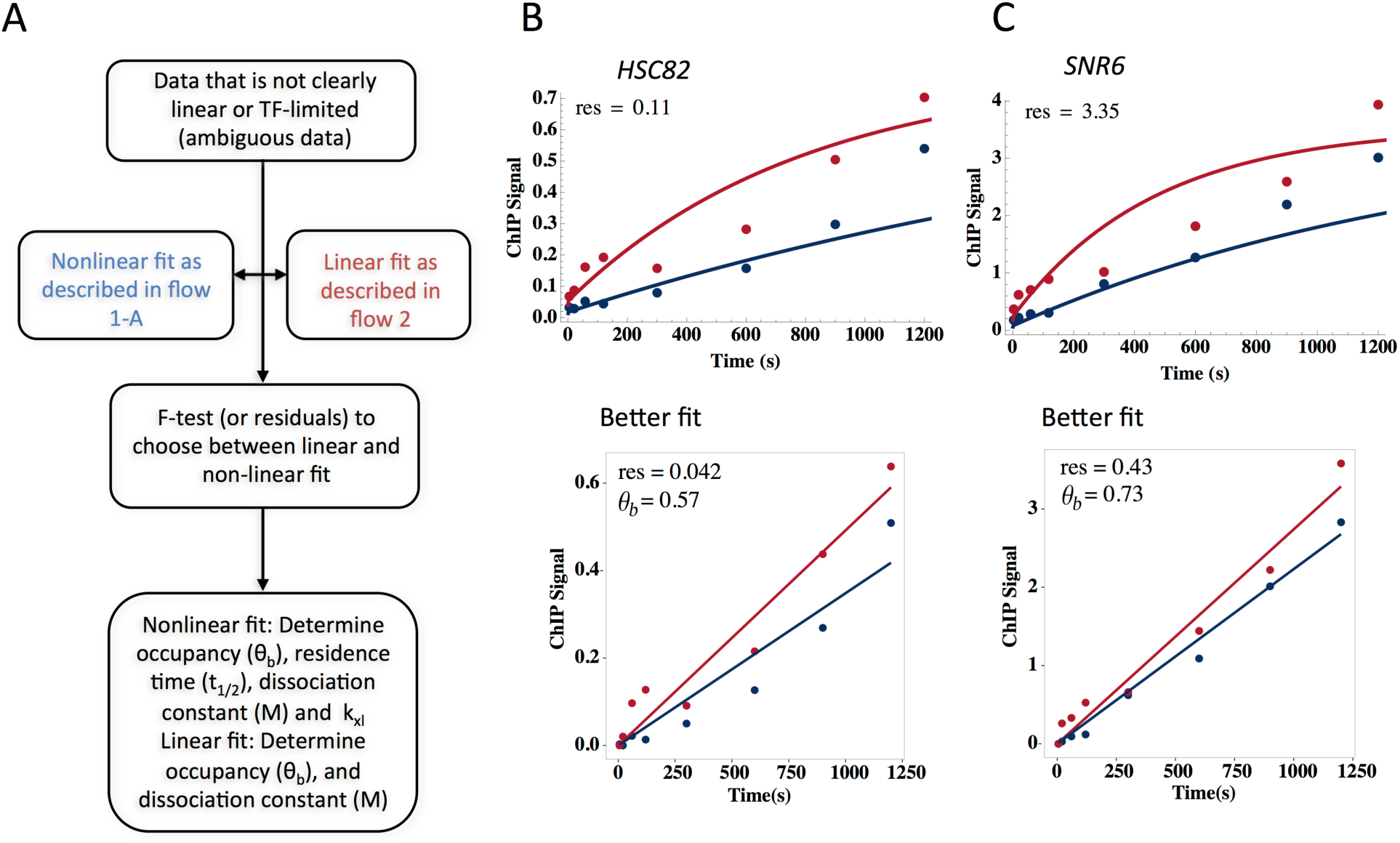
Resolution of ambiguous TBP fits. A) Flow chart to determine best fit for ambiguous data. Data was fit with both linear and full models and F-tests or sum of squared residuals (SSR) was then used to differentiate the best fit. B-C) TBP fits at *HSC82* (B) and *SNR6* (C) were fit with both full (top) and linear (bottom) models. SSR derived from the fits was used to find that both datasets were best represented with the linear fit; SSR is shown on all four plots and occupancy (θ_b_) for the linear fit.

As mentioned earlier, it is possible to model kinetic behavior of TFs that are depleted by formaldehyde by focusing measurements on the formaldehyde incubation time period where levels remain stable. TFIIE was significantly depleted by about ten minutes (Fig. 5A, E), but the protein levels were not detectably changed through seven minutes of formaldehyde incubation (Fig. 10A). This allowed us to measure TFIIE interaction with the *ACT1*, *LOS1*, and *URA1* promoters (Fig. 10B-D, Table 2). TFIIE binding to *URA1* and *LOS1* was best described by a crosslinking-limited model, whereas a full model fit described binding to *ACT1*. Fractional occupancies were well below saturation for all three sites, and at *ACT1* we compute a residence time of about 6 minutes, on par with the time-scale for TBP interaction at this site.

**Figure 10.**
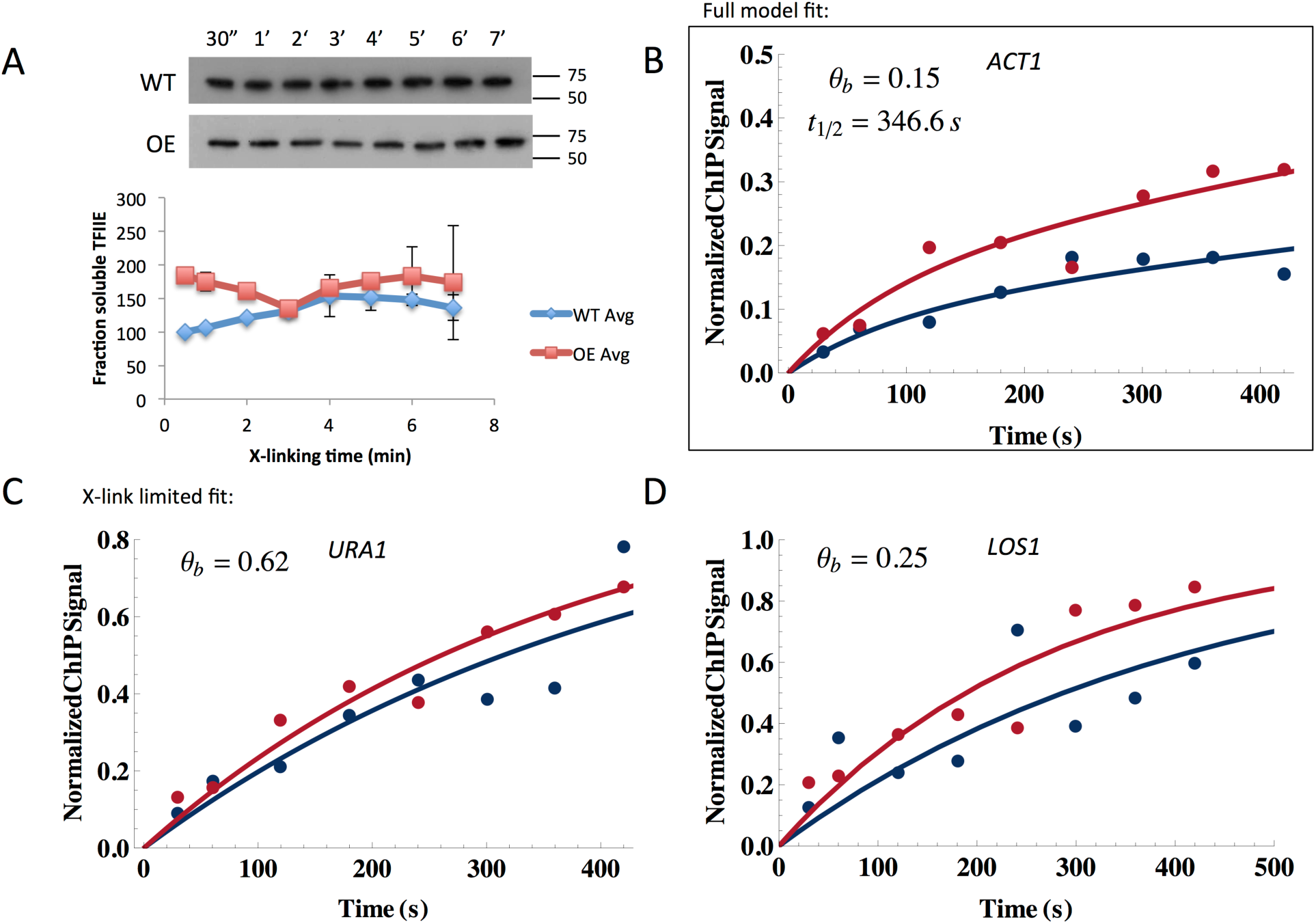
CLKv2 for TFIIE on a shorter experimental time scale. A) Western blot of Tfa1-TAP chromatin using an anti-TAP antibody. Samples were crosslinked for 30 seconds to seven minutes. Wild type and overexpression strains were both tested for depletion. Quantification of the signal was plotted below; two replicates were averaged and the standard deviation is shown as error bars. The wild type strain was normalized to its 30 second time point; the overexpression strain was normalized to its 30 second time point and multiplied by the overexpression factor (Table S3). The overexpression factor was determined by running four 5% formaldehyde-crosslinked time points for the wild type and overexpression strains on the same gel and blotting for TAP tag (data not shown). Bands were quantified with ImageJ (NIH) and compared to determine overexpression. B) TFIIE at *ACT1* resulted in a full model fit; occupancy and residence time are denoted. C) Both TFIIE at *URA1* and *LOS1* were crosslink-limited fits; only the occupancy is shown.

**Table 2.**
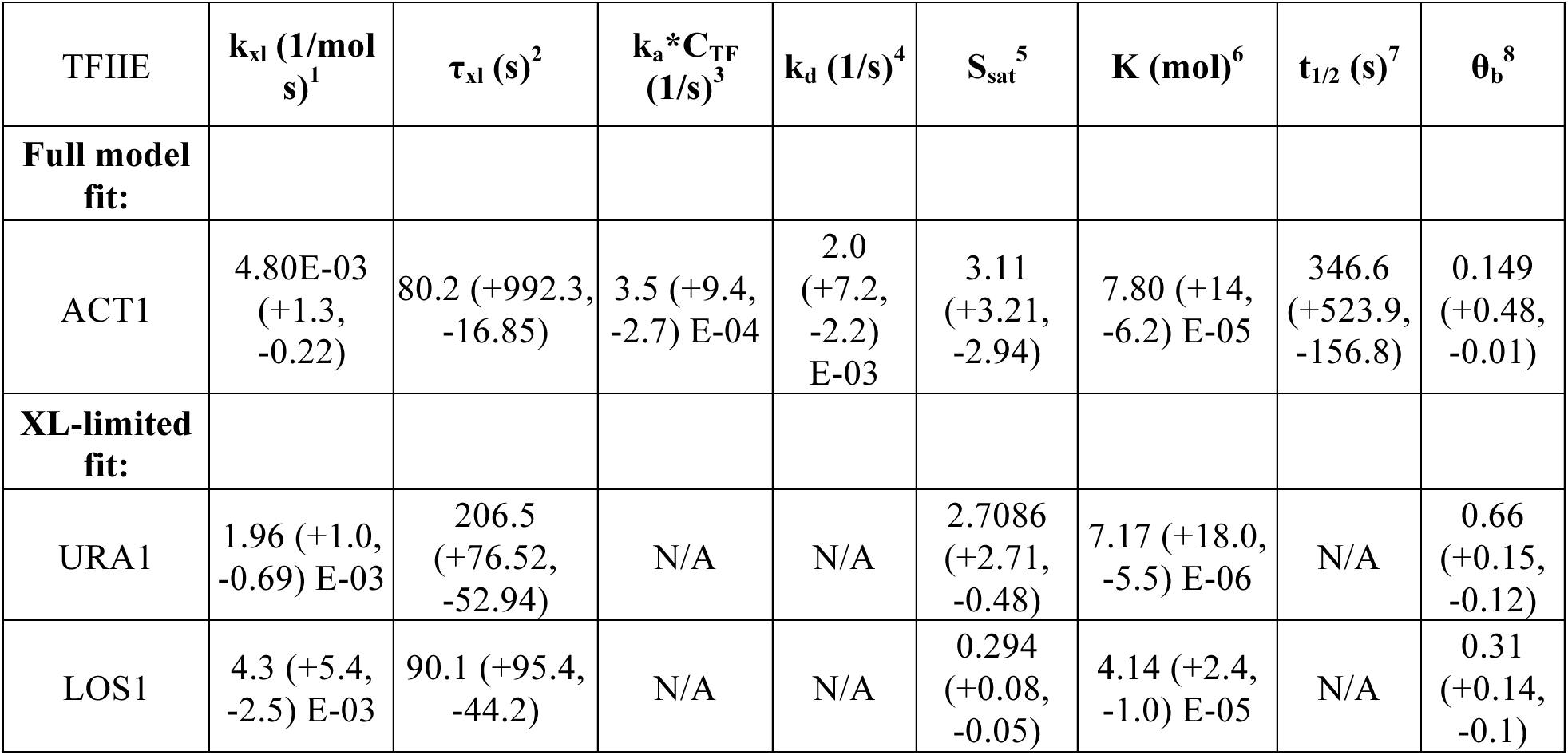
Measurements for TFIIE binding dynamics at select promoters.

## DISCUSSION

The CLK assay was conceived to provide biophysically rigorous on and off rates for TF binding to single copy loci in vivo (15). We sought to develop an approach that would also be generally applicable and potentially scalable to genome-wide analysis. The biggest obstacle to implementation of this assay has been to develop general experimental conditions and a companion model that accurately account for the many effects occurring in cells that undergo formaldehyde crosslinking and to distinguish them from the contributions of binding kinetics to the time dependent change in ChIP signal. Here we extend our understanding of the effects of formaldehyde on yeast cells and use our observations to both improve the CLK assay conditions and to improve the approach to data analysis. Formaldehyde crosslinking is ubiquitous in the chromatin field, so the results that we report here may contribute to the understanding and interpretation of ChIP and related types of experimental results in general as well.

Our results demonstrate improvement in formaldehyde quenching using a higher concentration of glycine than was used previously. The residual unquenched formaldehyde that remains following addition of 250 mM glycine as commonly used and in the original CLK procedure likely inflated the ChIP signal values at short crosslinking times as the unquenched formaldehyde continued to capture complexes during the centrifugation step that follows quenching. However, despite this, the relative differences in ChIP signal change with time apparent in the original CLK data do capture the relative differences in binding dynamics validated by other methods. For example, the rapid rise in Ace1 ChIP signal with short crosslinking times observed originally is consistent with the known highly dynamic behavior of Ace1 binding to its sites in the *CUP1* promoter (18), whereas the shallow slope and gradual approach to saturation seen with LacI time-dependent ChIP signals are consistent with its long residence time (19), which we confirmed by live cell imaging (15). Remarkably, the residence times for TBP binding to particular promoters reported here are also broadly consistent with the residence times obtained with the original version of the CLK assay (15). The results argue that TBP has residence times at these promoters on the order of one to several minute time scale. Thus, although the original CLK data was modeled assuming infinitely fast quenching, we nonetheless captured the relative time scale of dynamic behavior as validated by both live cell imaging and in this study using CLKv2.

Based on the results presented here, although Tris is highly effective in quenching unreacted formaldehyde, it is unsuited for use in this type of kinetic analysis due to its ability to reverse crosslinks. The crosslink reversal that we observed is consistent with a prior report (28) and is exacerbated by the relatively high concentration of Tris required to completely react with a relatively high concentration of added formaldehyde. We also show that time-dependent increases in ChIP signal can be affected by the concentration of formaldehyde. The use of a formaldehyde concentration that is as high as possible boosts the crosslinking rate, thereby extending the useful range of the assay. Although we employed 5% formaldehyde here, this may not be advisable or appropriate for analysis of other TFs or in other types of cells. The best formaldehyde concentration ought to be determined empirically by choosing the concentration that yields the best separation between the crosslinking and binding dynamics time scales, and which does not impact overall recovery of soluble components or deplete the unbound TF in the soluble pool over the kinetic time course. Those factors that are stable constituents of multi-subunit complexes such as Rpb1 may be impossible to assess using this approach; what is observed by western blotting as their rapid depletion from extracts may be due to rapid crosslinking to other biologically relevant polypeptides with which they stoichiometrically co-associate.

Using the CLKv2 method, we find that crosslinking rates are highly variable and depend on the particular TF-DNA site of interaction (Table 1). Prior to our measurement of formaldehyde crosslinking rates in vivo, crosslinking of ChIP complexes was generally thought to be rapid (14, 34, 35, 36), and this was supported qualitatively by the differences in ChIP signals that were observed at closely spaced time points (26) and that highly transient interactions (residence times on the ~second scale) could nonetheless be captured by formaldehyde crosslinking in ChIP experiments (8, 15, 16, 37). In addition, there is a global correlation between steady-state ChIP signals and in vitro binding affinity (38, 39) consistent with the overall ChIP signal level not being merely proportional to the rate of capture by crosslinking. In vitro, the rate of formaldehyde reaction with DNA bases is relatively slow (40), but reactivity could be greatly accelerated when DNA and amino acids were present together (41). Interestingly, the rates of TBP crosslinking to the *URA1* and *ACT1* promoters calculated by CLKv2 (*k_xl,_* Table 1) are in the same range as in vitro crosslinking rates obtained in reactions containing DNA and amino acids (41). Experiments measuring formaldehyde reactivity with amino acids and proteins have shown that formaldehyde adducts tend to be mainly formed with cysteine, lysine, and tryptophan side chains as well as the N-terminal group of polypeptide chains (42, 43). In reactions containing both nucleic acids and protein/amino acid substrates, the most efficient crosslinking was found to occur between lysine and deoxyguanosine (13, 14, 15). We suggest that the wide range in crosslinking rates reported here reflects the variation in reactive chemical groups on the TF surface and their proximity and orientation to reactive groups on DNA bases at or near binding sites.

Although some factors of interest were eventually depleted from extracts following formaldehyde treatment, our results with the TFIIE subunit Tfa1 show that it is still possible to investigate them kinetically if the crosslinking time course is confined to a temporal window in which their overall levels are not affected by formaldehyde. A possible limitation in this approach is that a shorter time course may make it more difficult to determine the saturation level of the ChIP signal, an estimate for which is required for confident fitting of the data and accurate estimates of the parameters. An alternative approach for future work is to extend the current model to include the depletion of the TF of interest in the fitting. Conceptually, by quantifying the rate of TF depletion from Westerns such as those shown in Fig. 5, the decrease in the overall level of the TF with crosslinking time could be modeled and the level of the TF at different times included explicitly as a parameter during the analysis of the data.

In instances in which the crosslinking rate is comparable to or slower than TF-DNA binding, CLKv2 yields the fractional occupancy as well as the equilibrium binding constant. Although the residence time cannot be estimated from the data in these situations, the fractional occupancy and binding constant are useful parameters as they provide insight into the variation in site occupancy across the cell population, which could have implications for understanding the molecular basis of transcriptional noise (44, 45), as well as energetic barriers in the intracellular environment that reduce binding from in vitro values obtained using purified components. If the crosslinking rate can be determined, this can be used to set an upper limit for binding dynamics. For many biological systems, knowing whether binding is occurring faster or equal to the second, minute or tens of minutes time scale would be valuable for developing dynamic models for the order of events underlying transcriptional responses.

## EXPERIMENTAL PROCEDURES

### Yeast Strains and Growth Conditions

Many of the *S. cerevisiae* strains used in this study were described previously (15); other strains were newly developed for the work presented here and all are listed in Table S1. TBP ChIP was performed in two ways: (1) using a monoclonal antibody that recognizes untagged TBP, and (2) using an antibody that recognizes the epitope tag on TBP-myc. Chromatin-associated myc-tagged TBP was measured using the epitope-tagged strain YAD154. TBP ChIP using the monoclonal TBP antibody was performed in various strains as described below. YAD154 cells used for the TBP-myc ChIP experiments were grown in YPD overnight at 30°C and harvested at OD_600_ ~ 1. For other TBP ChIP experiments comparing strains with two different levels of TBP, AY146 (wild type TBP levels) and YSC018 (harboring a 2μ TBP overexpression plasmid) were obtained from the TBP shuffling strain YAD165 as described previously (15). Cells were grown in synthetic medium without leucine plus 2% glucose overnight at 30°C. Culture volumes for each type of experiment are noted below and range from 100-450 ml depending on the experiment. When an OD_600_ of ~0.8 was reached, cells were pelleted and resuspended in an equivalent volume of YEP plus 2% glucose medium. They were grown at 30°C for approximately one hour until an OD_600_ of 1.0 was reached and cells were then formaldehyde crosslinked as described below. This regimen allowed cells to be initially grown under plasmid selection, but then transferred to YPD in order to standardize ChIP results which could otherwise be potentially influenced by effects of growth medium, and in addition, growth in YPD prior to crosslinking permitted direct comparison with previously published work (2, 3).

For TFIIE ChIP, strains were used with WT or elevated levels of the two TFIIE subunits Tfa1 and Tfa2 in which the Tfa1 subunit was TAP tagged. YRV006 (*TFA1*-TAP, Dharmacon) (15) was transformed with an empty pRS315 vector (46) (AY151, WT TFIIE levels) or a pRS315- based plasmid carrying copies of *TFA1*-TAP and *TFA2* under control of their endogenous promoters (AY152, over-expressed levels of TFIIE). Cells were grown at 30°C overnight in synthetic medium without leucine and with 2% glucose. When an OD_600_ of ~0.8 was reached, cells were pelleted, resuspended in an equivalent volume of YPD, and grown at 30°C to an OD_600_ of 1.0 as described above. Strain construction for AceI (YTK539 and YSC002) and LacI (YTK260 and YSC001) as well as collection for ChIP was described previously (15).

For western blotting, strains YGR186W, YBR049C, YRV018, and ML307-1 were grown overnight in YPD at 30°C to _OD600_ of 1; YRV005 was grown in YEP + 2% raffinose at 30°C overnight to OD_600_ 0.8, then 2% galactose was added and cells incubated to OD_600_~1.0. YTK539 cells were grown under conditions of copper induction as previously described (15).

### Quenching and crosslinking conditions

Different crosslinking and quenching conditions were tested with the TBP-myc strain (YAD154) in order to explore the relationship between crosslinking rate and formaldehyde concentration, as well as quenching efficiency. In all experiments, cells were first grown in YPD at 30°C overnight to an OD_600_ of 1.0. To test the effect of 250 mM glycine, 100 ml cell cultures were incubated with 2.7 ml 37% formaldehyde (1% final, Fisher) followed by addition of 10 ml 2.5 M glycine (pH 6.3) at various times. To test the effect of 2.93 M glycine, 450 ml cultures were grown in YPD overnight at 30°C to OD_600_ of 1. Cells were then concentrated five-fold by centrifugation and resuspended in 90 ml YPD. The concentrated cultures were then incubated with 2.7 ml 37% formaldehyde (1% final concentration) by addition of formaldehyde to the culture while rapidly mixed using a stir bar. At various times thereafter, 10 ml aliquots were removed and added to 440 mL glycine pH 5 contained in 450 ml Sorvall centrifuge bottles. Bottles were capped by hand as quickly as possible and vigorously shaken. Samples were washed and worked up as detailed below.

To test different formaldehyde concentrations and other quenching conditions, TBP-myc cells were grown as described above. To test formaldehyde concentrations at 1% or lower, in most cases the appropriate volume of 37% formaldehyde was added to a rapidly stirring 100 ml culture, and the reaction was then quenched after specific incubation times by addition of 3 M glycine or 3 M Tris-HCl, pH 8, to achieve the indicated final quencher concentration. For reactions in which formaldehyde was added to a final concentration greater than 1%, cells were concentrated five-fold in YPD as described above, 37% formaldehyde was added to achieve the indicated final concentration, and after particular incubation times, 10 ml aliquots were removed to centrifuge bottles or tubes containing 3 M glycine or Tris yielding the final concentration of the quencher indicated in the figure legends. Cell samples quenched in Tris were worked up and analyzed as described above except that the first TBS wash contained 120 mM Tris-HCl pH 8 rather than glycine.

### Quenching reversal experiments

To determine the stability of crosslinked material in the presence of quencher, crosslinked cells were incubated in solution containing glycine or Tris for different periods of time prior to ChIP work-up. For Tris-quenched samples, 100 ml cultures of AY146 cells were grown overnight in synthetic media lacking leucine and containing 2% glucose at 30°C. Cells were then transferred to YPD at an OD_600_ of 0.8 and grown until reaching an OD_600_ of 1. Each sample was crosslinked by adding formaldehyde to 1% for 5 minutes and then quenched by adding 10 ml 2.5 M glycine to each 100ml culture. Cells were pelleted and resuspended in either 750 mM Tris-HCl pH 8, or TBS buffer (which contains 50 mM Tris-HCl, pH 8 as described above) and incubated at room temperature for 10 or 30 minutes. Subsequent steps were carried out as described below.

To test crosslink stability in the presence of glycine, 250 ml replicate cultures of AY146 cells in synthetic media plus 2% glucose and without leucine were incubated overnight at 30°C, then resuspended in YPD and grown to an OD_600_ of 1 as described above. Three aliquots of 50 ml were taken from each culture and pelleted at room temperature. Each pellet was then resuspended in 10 ml YPD and transferred to a flask on a stir plate. Formaldehyde was then added to 5% final concentration to each sample and mixed at room temperature for 5 minutes. 10 ml from each sample were quenched in 440 ml 3 M glycine pH 5 at room temp for 0, 10, or 30 minutes. The zero minute sample was pelleted at 4°C immediately after quenching; the other time point samples were pelleted the same way after glycine incubation of 10 or 30 minutes. Following incubation of the crosslinked cells in glycine solution for the indicated times, the cells were processed for ChIP as described below.

### Order-of-addition experiments

Order-of-addition experiments were performed to test quenching efficiency using the TBP-myc strain, YAD154. Replicate cultures of YAD154 cells (300 ml) were grown overnight at 30°C in YPD to an OD_600_ of 1.0, then concentrated by resuspension in 60 mL YPD. In each experiment, three 10 ml aliquots were collected in duplicate: (1) no formaldehyde control samples in which 3 M glycine pH 5 was added to 2.93 M final concentration, (2) samples in which 3 M glycine was added to 2.93 M final concentration before 5% formaldehyde addition for 8 minutes, and (3) 5% formaldehyde incubation for 8 minutes followed by addition of 3 M glycine pH 5 to 2.93 M final concentration. Following these treatments, cell samples were washed in 50 ml TBS plus 300 mM glycine pH 5 followed by washing in 50 ml TBS, both washes at 4o C. Subsequent work-up for ChIP and Real Time PCR for TBP binding to the *URA1* locus were performed as described above.

Order-of-addition experiments for Gal4 with the previously published CLK conditions (15) were done in the same way as order-of-addition experiments described above, except different glycine and formaldehyde concentrations were used. For each sample set, three 100 ml YPH499 cultures were grown overnight at 30°C in YEP + 2% raffinose. When an OD_600_ of 0.8 was reached, each culture was induced with 2% galactose. At OD_600_ of 1.0, samples were collected in duplicate. The following experimental parameters were used: (1) 2.5 M glycine pH 6.3 was added to 250 mM final concentration, (2) 2.5 M glycine pH 6.3 was added to 250 mM final concentration for 5 minutes before addition of 1% formaldehyde for 8 minutes, and (3) 1% formaldehyde incubation for 8 minutes before addition of 2.5 M glycine pH 6.3 to 250 mM final concentration for a 5 minutes incubation. The subsequent steps were the same as above, except analysis was performed for interaction at the *GAL3* locus.

### Whole cell extract preparation and western blotting

Strains were grown in 300 ml YPD overnight to an OD_600_ =1.0 in YPD, then concentrated five-fold as described above. Following removal of a zero minute (no formaldehyde) control, formaldehyde was added to 5% and cells were incubated for various times at room temperature as indicated in the figures and then 10 ml aliquots were quenched in 440 ml of 3 M glycine pH 5. Samples were spun down and then prepared as either chromatin or whole cell extracts (WCE); the Benoit’s buffer (200 mM Tris-HCl (pH 8.0), 400 mM (NH_4_)_2_SO_4_, 10 mM MgCl_2_, 1 mM EDTA, 10% glycerol, 7 mM bmercaptoethanol) lysis extraction protocol was employed for whole cell extracts (2, 3). Chromatin extracts for western blotting were prepared in the same way as chromatin was prepared for ChIP. The only difference for chromatin samples was the use of 300 mM glycine pH 5 in the first TBS wash instead of 250 mM glycine pH ~6.3 (2, 3). Both chromatin and WCE protein levels were quantified with Bradford protein dye (Bio-Rad) using bovine serum albumin as the standard. 8% or 10% denaturing protein gels were used to resolve 15 μg protein for each sample. Unless otherwise noted, before loading the gel, samples were incubated at 95°C for five minutes. This heating step was left out for unheated samples. Coomassie staining or membrane transfer was performed following electrophoresis. For staining, the gel was incubated with coomassie dye (Research Organics Inc) for one hour at room temperature with gentle shaking, followed by overnight destaining (40% methanol, 10% acetic acid) at room temperature. The gel was imaged with the FluorChemQ system (protein simple). For gel transfer, proteins were transferred to Immobilon P and western detection of particular protein species was performed using the antibodies listed in Table S5 and detection with Amersham ECL Prime (GE Healthcare). Quantification of bands on the blots was done using ImageJ software (NIH).

### Collection of crosslinking time points and preparation of chromatin samples

We found that collection of eight crosslinking time points in a single experiment was manageable. A single eight time point experiment performed with optimized glycine quenching required nearly 4 liters of 3M glycine, which was made by adding 900.84 g glycine (Bio-Rad) to a total volume of 4 L water. The solution was gently heated on a hot plate to help the glycine dissolve. The pH of the resulting solution was then adjusted to 5 using a few milliliters of concentrated HCl (Fisher). The glycine was then aliquoted into eight 500 ml bottles, each of which contained 440 ml of the solution. The flask containing 90 ml cell culture was rapidly mixed with a stir bar, and 14 ml 37% formaldehyde (Fisher) was added to the culture (resulting in 5% final formaldehyde concentration) at time zero. 10 ml aliquots of culture were then removed from the flask using a Pipet Aid and immediately added to the aliquoted glycine solution. For each sample, bottles were immediately capped and vigorously shaken for a few seconds to ensure good mixing. All subsequent steps were performed at 4°C by keeping the samples on ice, and using buffers and centrifuges chilled to 4°C. Quenched cell samples were pelleted by centrifugation for 7 minutes at 5000 rpm in an SLA-3000 rotor. Cell pellets were resuspended in 50 ml TBS plus 300mM glycine and transferred to 50 ml conical tubes. The tubes were centrifuged for 5 minutes at 4000 rpm in a clinical centrifuge. Cell pellets were then washed with 50 ml TBS (40 mM Tris-HCl pH 7.5, 300 mM NaCl) and spun as before. Each pellet transferred to a FastPrep tube and cell pellets were stored at -80°C for later work-up or resuspended in 600 μl 140 mM ChIP lysis buffer (50 mM Hepes pH 7.5, 140 mM NaCl, 1% Triton X-100, 0.1% sodium deoxycholate) with protease inhibitors (Roche Complete Protease Inhibitor Cocktail Tablet OR 1.0 mM phenylmethylsulfonyl fluoride, 2.0 mM benzamidine, 2.0 mM pepstatin, 0.6 mM leupeptin, and 2.0 mg of chymostatin per ml of buffer) for bead beading.

Once pellets were resuspended in ChIP lysis buffer, acid-washed beads (Sigma) were added to just above the liquid line and samples were processed for 7 cycles of 45 sec on, 1 min off in a FastPrep machine (MP Biomedicals). Tube bottoms were punctured with an 18-guage needle (BD PrecisionGlide) and placed in 13 x 100 mm glass tubes and the liquid recovered by centrifugation for 5 min at 3000 rpm. Each sample was briefly vortexed and then transferred to a 1.5 ml eppendorf tube on ice. Samples were then sonicated with a Branson Sonifier 250 with microtip probe for 7 cycles of 5 pulses each with 30% output and 90% duty cycle. This was followed with a 5 min spin at 14000 rpm and 4°C. The supernatant was transferred to a new eppendorf tube. Following a second spin for 20 min at 4000 rpm, supernatants were collected and the protein was quantified by Bradford protein assay as described above.

### ChIP and real time PCR

Chromatin immunoprecipitation was performed with 1 mg total protein for each sample. For each time point IP, mock, and total (input) samples were assayed. IP and mock sample volumes were adjusted to 500 μl with 140 mM ChIP lysis buffer with protease inhibitors added. For TBP ChIP, 2.5 μl of anti-TBP antibody (Cat# ab61411, Abcam) was used in the IP. For TBP Myc, 2.5 μl of anti-Myc antibody (Cat#ab32, Abcam) was used. For LacI and AceI, 5 μl of anti-GFP antibody (Cat# A11122 Life Technologies Inc) was added to samples. The IP and mock samples were inverted overnight at 4°C. Following overnight incubation, the IP and mock samples were then incubated with 40 μl Sepharose A Fast Flow 4 beads (GE Healthcare) for 2 hours at 4°C. Samples were washed twice with 1 ml of 140 mM ChIP lysis buffer, 500 mM ChIP lysis buffer (same as 140 mM ChIP lysis buffer but containing 500 mM NaCl), LiCl wash buffer (10 mM Tris pH 8.0, 250 mM LiCl, 0.5% NP-40, 0.5% sodium deoxycholate, 1 mM EDTA), and 1X TE (10 mM Tris-Cl pH 8.0, 1 mM EDTA). Two elutions of the bound material were performed by adding 75 μl elution buffer (50 mM Tris-HCl pH 8.0, 1% SDS, 10 mM EDTA) to each sample for 10 min at 65°C. The two elutes were combined and incubated overnight at 65°C along with the total samples, which consisted of 0.1 mg input chromatin protein combined with 150 μl elution buffer. The following day, samples were cleaned up using the QiaQuick PCR cleanup kit (Qiagen) following the manufacturer’s instructions and DNA was eluted with 50 μl DEPC water pre-warmed at 55°C.

ChIP for TFA1-TAP was performed as described above, except 40 μl of a 50% slurry of IgG Sepharose 6 Fast Flow beads (GE Healthcare) was added to the IP sample and 40 μl of a 50% slurry of Sepharose 6 Fast Flow beads (GE Healthcare) was used for the mock samples. An overnight IP was carried out at 4°C followed by washing the bead pellet the next day as described above.

To quantify the ChIP DNA, real time PCR was performed using appropriate primer sets, iQ SYBR Green Supermix and a MyiQ instrument (Bio-Rad). The standard curve inputs were run in duplicate and all unknowns (IP, mock, total samples) were run in triplicate. The relative ChIP signal for each time point was calculated by subtracting the mock signal from the IP signal and then dividing by the total signal. The kinetic data reported here represent the average from at least two independent experiments for each strain and condition.

### Computational Modeling

The Crosslinking Kinetics (CLK) model is described by Eqns. 7, 11, and 16 in Sec. 2.2 of Poorey et al. (15) Supplementary Material. The model is characterized by the transcription factor association rate (k_a_) and disassociation rate (k_d_) of binding to chromatin, the formaldehyde-transcription factor crosslinking rate (k_xl_), the saturation level of the ChIP signal (S_sat_), the transcription factor concentration *in vivo* (C_TF_), and the formaldehyde concentration (C_FH_). The ChIP signal, S(t), is related to the *in vivo* fraction of a given binding site cross-linked by the TF across cells (θ_xl_) by the relationship θ_xl_(t) = S(t)/S_sat_ where S_sat_ is the saturation value of the ChIP signal. This scaling of the ChIP signal ensures that θ_xl_(t) approaches 1 as crosslinking time goes to infinity, as required by the CLK model. Two physically interpretable parameter regimes of the model are the transcription-factor dynamics limited (TF-limited) regime where TF dynamics are much slower than crosslinking dynamics (i.e., k_a_*C_TF_ << k_xl_*C_FH_ and k_d_ << k_xl_*C_FH_) and the crosslinking dynamics limited (XL-limited) regime where crosslinking dynamics are much slower than TF dynamics (i.e., k_xl_*C_FH_ << k_a_*C_TF_ and k_xl_*C_FH_ << k_d_), as detailed in Sec. 2.3 of Poorey et al. (15) Supplementary Material. Finally, for extremely slow crosslinking dynamics which occur on the timescale of the full range of crosslinking times or longer (i.e., k_xl_*C_FH_*θ_b_*t_l_ << 1 where t_l_ is the last crosslinking time point, which is usually 1200s), the CLK model predicts that θ_xl_(t) will be a linear function of crosslinking time, θ_xl_ ~ k_xl_*C_FH_*θ_b_*t. Notably, we observe TF-limited, XL-limited and linear in crosslinking time CLK curves depending on the TF and locus examined.

The simulations presented in Fig. 6 show the expected CLK curves in the TF-limited, the XL-limited, and the linear regimes, while the schematic diagram shows the physical interpretation of the *in vivo* dynamics in these regimes. The hallmarks of the TF-limited model are a relatively fast exponential rise at time scales of less than ~100 seconds but often less than 5 seconds (first crosslinking time point in the experiment) followed by a slower exponential rise (see Fig. 6A). Notably, when the first relatively fast exponential rise is less than 5 seconds, we observe a non-zero y-intercept in the WT and OE data with a clear separation between the WT and OE y-intercepts. When the rise in the first relatively fast exponential is ~100 seconds, we find a zero y-intercept, an initial fast exponential rise in the data followed by a slower exponential rise, hence forming what looks like a “knee” in the data around the transition from the fast to the slow exponential for both the WT and OE data. Interestingly, the y-intercept for very fast crosslinking or “knee” for modestly fast crosslinking in the WT data yields an excellent approximation of the in vivo occupancy, θ_b_. The XL-limited model shows a single exponential rise with a zero y-intercept for the WT and OE data (see Fig. 6B). The linear model shows a near-zero y-intercept at t=5 seconds, and no sign of saturation on the experimental time scale of 700 seconds to 1200 seconds (see Fig, 6C). Importantly, the two crosslinking dynamics limited models, XL-limited and linear, display relatively high sensitivity to formaldehyde concentration (as shown in Fig. 1B,C) while the TF-limited (which we also refer to as the “full model” for reasons described below) does not. While the full mathematical model presented in equations (11) and (16) in the Supplemental Material of Pooery et al (15) can be used to fit and represent all of these parameter regimes, we use and refer to a “full model” fit for data that clearly show the double exponential behavior (i.e., relatively fast crosslinking rise followed by a second TF-dynamics limited rise with a relatively clear kink or knee in between the two). Moreover, in the case of XL-limited behavior, we use the single exponential XL-limited model shown in equation (21) of the Supplementary Materials of Poorey et al (15), which is a highly accurate approximation of the “full” model (equations (11) and (16) in the Supplemental Material of Poorey et al (15)) in the XL-limited parameter regime. Finally, for linear in crosslinking time data, we use the linear model shown in equation (22) of the Supplementary Materials of Poorey et al (15), which is a highly accurate approximation of the “full” model in the very slow crosslinking dynamics parameter regime.

For data that showed negative curvature (i.e., TF-limited or XL-limited), we started by visually estimating S_sat_ to be close to the late time point over-expression ChIP signal. Hence, our initial guess was normally S_sat_ between 1 and 5, except for LacI, where we started with S_sat_ ~10. In the case of data that visually showed TF-limited behavior (e.g., TBP at ACT1, LOS1, and URA1), we estimated the initial value for k_xl_ by looking at the time (τ_xl_) around which the data showed a “knee.” Setting ln[2]/k_xl_ ~ τ_xl_ gives an estimate for k_xl_. The y-intercept of a linear extrapolation of the late-time S(t) data points (i.e. linear extrapolation of the S(t) data points that are approximated by the second exponential) divided by S_sat_ gives an initial estimate for θ_b_. The in vivo occupancy, θ_b_, is expressed in terms of k_a_ and k_d_ as θ_b_=k_a_*C_TF_/(k_a_*C_TF_+k_d_). For a given θ_b_, we can sweep over a wide range of k_a_ and S_sat_ values to see where the theoretical curves match with the WT and OE experimental data. Importantly, the overall on-rate, k_a_*C_TF,_ dominates the rate at which the second exponential rises. With these starting estimates for the kinetic parameters, we run the NonLinearModelFit routine in Mathematica (47) to fit the full model to the data using least squares. The fit reliably gives us k_a_, k_d_, and S_sat_ (equivalently, S_sat,_ θ_b_ = k_a_*C_TF_/(k_a_*C_TF_+k_d_), and t_1/2_=ln(2)/ k_d_).

For data that did not show TF-limited behavior (but still showed negative curvature, as opposed to a purely linear response, for example, TFIIE at ACT1 and URA1), there were two possibilities: either the data was XL-limited (showing a single exponential), or the knee was not markedly visible by inspection because of the experimental time scales. We started by fitting a straight line to the short crosslinking time data to estimate k_xl_*S_sat_ and θ_b_. With these estimates, we swept over a wide range of k_a_, k_xl,_ and S_sat_ values to match the theoretical full model with the data. With these tuned estimates, we fit both the XL-limited model and the full model to the data, and determined which model yielded a better fit of the data by looking at the validity of parameters obtained, the sum of squared residuals (SSR), or by conducting an F test.

For data that fit the linear XL-limited model best (e.g., TBP at NTS2 and ACE1), we subtracted the y-intercept (extrapolated ChIP signal at t = 0 second) from the data as background, and fit a line to each of the WT and OE data using least squares. The overexpression factor is known, so we could extract k_xl_*S_sat_ and θ_b_ from the two slopes.

For some loci it was not obvious if the data would fit the full model/TF-limited model or the linear XL-limited model (e.g., TBP at HSC82 and U6). It was important to answer the question of the better fit because the two models have a different number of effective parameters: the full-model fit has four free parameters (S_sat_, k_a_, k_d_, and k_xl_), while the linear regime has only one: S_sat_*k_xl_*θ_b_. The sum of squared residuals (SSR) with the linear fit (with fewer degrees of freedom) was lower than the SSR with the full model fit (with more degrees of freedom); hence, the linear fit was chosen without the need to conduct an F-test comparing the two models. The full model fit gave worse SSR values because we were explicitly starting with estimates close to the TF-limited regime when fitting the full model, which lead the minimization of the difference between the model and data to a suboptimal, local minimum. Note that the SSR was calculated without normalizing the data using S_sat_ because the SSR scales with Ssat and S_sat_ is unknown in the linear fit case.

For TFIIE at *ACT1*, the final parameters from the XL-limited fit were unphysical (θ_b_ ~ 0 and K_d_ = k_d_/k_a_ ~ 10^7^ mol); hence, the full model fit was chosen. An F-test was performed to choose the XL-limited fit for TFIIE at *LOS1* over the full model fit. For TFIIE at *URA1*, the parameter estimates from a full model fit satisfied XL-limited binding dynamic conditions. Therefore, the TFIIE data at *LOS1* and *URA1* were fit with the XL-limited model.

To estimate the errors associated with our output parameters, we ran our fitting procedure on simulated data for each locus. Specifically, we simulated the data at each locus with the mean value at each time point given by the theoretical fit and the variance given by the mean of the squared residuals. We simulated and fit the data at each locus for one thousand successful fitting iterations. The standard deviation in the simulated fit parameters was calculated on the log scale, and was transformed back from the log scale to determine the lower and upper bounds on the error bars quoted in Table 1. Error bars for k_xl_ could not be estimated in the case of TBP at *LOS1* since the fit parameters were TF-limited and fitting the full-model to the simulated data gave spurious values for k,d in addition to failing often. Hence the error bars for k_a_, CTF, k_d_ and S_sat_ for TBP at *LOS1* were calculated by fitting the TF-limited model to the simulated *LOS1* data.

## Acknowledgements

We are grateful to Nicole Francis for discussions and for her input on development of the method. This research was supported by NIH grant R01 GM55763 (to D.T.A), NIH grant R21 GM110380 (to S.B and D.T.A), and NCI Cancer Training Grant T32 CA009109-38 and Wagner Fellowship (to E.A.H.). The content is solely the responsibility of the authors and does not necessarily represent the official views of the National Institutes of Health.

## Author Contributions

D.T.A. designed, performed, and analyzed the data for Fig 1. S.J.S. performed and analyzed the data for Fig 2, 3, 4 and 5 and collected data for Fig 8 and 9. E.A.H. performed and analyzed data for Fig 5, 10, and S1. H.Z. and S.B. developed the model and analyzed data in Fig 8, 9, 10, S2, and S3. H.Z. and E.A.H. made the figures. D.T.A., E.A.H., H.Z., and S.B. wrote the manuscript. All authors reviewed the results and approved the final version of the manuscript.

## FOOTNOTES

1 Unpublished data.

The abbreviations used are: PIC, *preinitiation complex;* TFs, transcription factors; CLK, *crosslinking kinetic assay;* TBP, *TATA binding protein*; OE, *over-expression*; k_a_, *on-rate*; k_d_, *off rate*; k_xl_, *formaldehyde crosslinking rate*; θ_b_*, fractional occupancy*; t_1/2_, *residence time*; S_sat,_ *saturation level of the ChIP signal*; C_TF_, *concentration of the TF in the nucleus*; C_FH_, *formaldehyde concentration*; SSR, *sum of squared residuals*; WCE, *whole cell extract*.

